# GenerRNA: A generative pre-trained language model for *de novo* RNA design

**DOI:** 10.1101/2024.02.01.578496

**Authors:** Yichong Zhao, Kenta Oono, Hiroki Takizawa, Masaaki Kotera

## Abstract

The design of RNA plays a crucial role in developing RNA vaccines, nucleic acid therapeutics, and innovative biotechnological tools. Nevertheless, existing techniques lack versatility across various tasks and frequently suffer from a deficiency of automated generation. Inspired by the remarkable success of Large Language Models (LLMs) in the realm of protein and molecule design, we present GenerRNA, the first large-scale pre-trained model for RNA generation, aiming to further automate RNA design. Our approach eliminates the need for secondary structure or other prior knowledge and is capable of *de novo* generation of RNA with stable secondary structures while ensuring its distinctiveness from existing sequences. This widens our exploration of RNA space, thereby enriching our understanding of RNA structures and functions. Moreover, GenerRNA is fine-tunable on smaller, more specialized datasets for particular subtasks. This flexibility and versatility enables the generation of RNAs with desired specific functionalities or properties. Upon fine-tuning GenerRNA, we successfully generated novel RNA sequences exhibiting high affinity for target proteins. GenerRNA is freely available at the following repository: https://github.com/pfnet-research/GenerRNA

## 1 Introduction

The design and engineering of RNA, molecules characterized by a chain-like structure comprised of ribonucleotides, are instrumental in the progress of advanced therapeutics and biotechnologies. Historically, RNA design methodologies have been grounded in empirical approaches and directed evolution in wet-lab settings, which is a process plagued by high costs and low efficiency[1]. Subsequently, the advent of computational techniques permitted the quest of RNA sequences with specific secondary structure, which further advanced RNA design[2]. However, the reliance on predefined RNA structural configuration and other a priori knowledge has imposed constraints on the adaptability of these methods. [3]. Today, the achievements of deep generative language models present new possibilities for further automation in RNA generation.

Recently, deep generative language models have made notable strides in the field of natural language processing. These models learn from massive unlabeled datasets, produce meaningful representations, and generate high-quality text[4]. This success has resonated in the biological and chemical domains[5][6], offering a new paradigm where the structure and function of biomolecules can be handled within the framework of a natural language model. For instance, ProGen, a large-scale protein language model, generates protein sequences with predictable functions across diverse protein families, also aiding in the exploration of protein space[7]. Similarly, in the realm of small molecule design, there has been a proliferation of published generative language models utilizing string-based molecule representations, garnering considerable attention[8]. As for RNA research, while pre-trained language models have demonstrated efficacy in RNA function and structure prediction[9], the generation of RNA sequences remains largely unexplored.

We present GenerRNA, a generative RNA language model built upon the Transformer decoder architecture[10]. Typically, Transformer models undergo pre-training on large text datasets, enabling them to learn the underlying syntax and patterns of language in an unsupervised manner. This process entails predicting subsequent words or characters in a text sequence without any reliance on labels or annotations. GenerRNA was pre-trained on approximately 30 million RNA sequences encompassing 17.4 billion nucleotides to acquire a broad, cross-family understanding of RNA representations, facilitating *de novo* sequence generation. We evaluated the generated RNA sequences from the following standpoints: secondary structural stability measured by Minimum Free Energy and novelty assessment by conducting homology searches. Additionally, the nucleotide propensities of the generated sequences are similar to those found in natural sequences. The results reveal that GenerRNA can generate RNA sequences that are novel, nature-like, and structurally meaningful. Furthermore, we fine-tuned the model to generate RNAs with the capability to bind to specific proteins. *In silico* evaluations indicated that these generated RNAs exhibit high affinity scores with target proteins. Additionally, ablation experiments demonstrated the substantial role of pre-training, allowing for the generation of more rational and novel sequences.

GenerRNA signifies a considerable advancement in automating RNA design. This innovation underscores the potential of generative language models as a promising and effective approach in RNA engineering.

## 2 Methods

### 2.1 Learning the RNA Language

GenerRNA is a language model that processes RNA sequences through a linguistic perspective. By employing unsupervised learning on a large-scale RNA dataset to discern the inherent syntax, grammar, and semantics in RNA sequences, thereby mastering the ability to “speak” the RNA language—signifying its capacity to generate RNA sequences akin to nature.

Within this framework, GenerRNA computes the probability for each specific token *x*_*i*_ in a sequence, where a *token* refers to a discrete unit consisting of one or more nucleotides. This probability is influenced by the token’s context, determined as the joint probability of preceding tokens in the sequence ranging from *x*_1_ to *x*_*i −*1_. The aggregate probability of a sequence *X* is derived from the joint probability distribution of its constituent tokens.

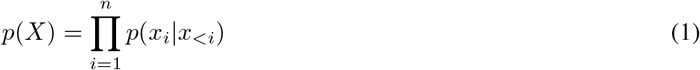

Focusing on sequence generation, GenerRNA employs an autoregressive approach, implying that the model operates in a sequential, left-to-right fashion to construct the sequence one token at a time. This process necessitates that the prediction of each subsequent token depends on all previously generated tokens, ensuring a contextually coherent progression in the sequence generation.

GenerRNA is a neural network based on the Transformer decoder architecture[10]. The Transformer architecture features a series of stacked layers designed to learn the global context within sequences, with each layer incorporating a self-attention mechanism as illustrated in Fig.1**b**. This self-attention mechanism within each layer is responsible for interpreting pairwise interactions among all positions in its input sequence. Our decoder-only architecture model consists of 350 million parameters across 24 transformer layers and has a model dimension of 1280. This configuration mirrors the GPT2-medium model architecture proposed by OpenAI[15].

**Figure 1:**
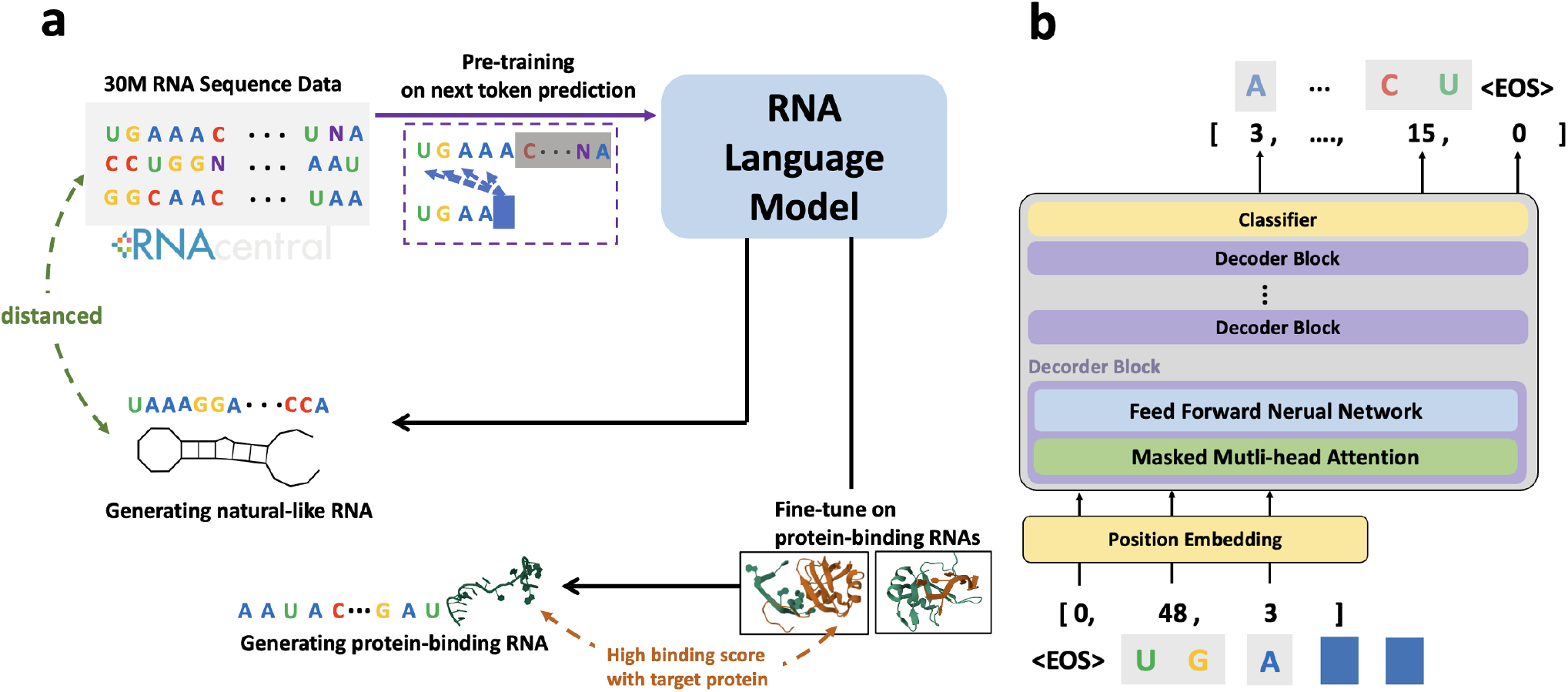
Overview of GenerRNA workflow and design architecture. **a**. GenerRNA undergoes unsupervised pre-training on a large-scale RNA corpus, enabling *de novo* sequence generation. The model can be further fine-tuned to perform specific downstream tasks. **b**. GenerRNA is composed of 24 Transformer decoder layers. The model operates in an autoregressive manner to predict the subsequent token. Both the input and output of the model are in the form of tokens, which are encoded and decoded by a trained tokenizer. The 3D structures in the diagram were sourced from RNAComposer[11] and the Protein Data Bank[12][13][14].

We trained the model by minimizing the negative log-likelihood (NLL) across the entire dataset. This approach involves the model learning the relationship of each nucleotide token in a sequence with the preceding nucleotides.

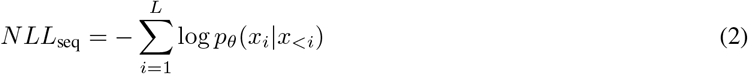

GenerRNA was trained on 18GB of sequence data, undergoing 20 epochs on a global batch size of 128, which equates to approximately 330,000 iterations.

The parameter weights were initialized prior to training. The optimizer used was Adam (*β*_1_=0.9, *β*_2_=0.999)[16] with the learning rate experiencing a warm-up over the first 10,000 iterations to 1e-3 with a linear decay to 1e-4 for the remainder of the training. The training was executed on 16 A100 GPUs and was completed within a week.

### 2.2 Pre-training Data Preparation

For pre-training, we obtained a sum of 34.39 million sequences from RNAcentral (release 22), a comprehensive database that combines RNA sequences from 51 expert databases[17]. The training data comprised over 2,600 Rfam families and 30 types of RNA types such as rRNA, tRNA, and lncRNA, but notably excluded mRNA [18]. In our preprocessing steps, we replaced ‘U’ for ‘T’ while retaining all ambiguity codes, and deduplication was accomplished using CD-HIT[19] with an identity threshold of 100%. Consequently, we selected sequences shorter than 256 tokens once tokenized. This selection criterion resulted in a refined dataset containing 29.75 million sequences (18.15GB), encompassing 17.4 billion nucleotides. 99% of the filtered sequences were allotted for training, while the remaining 1% composed the validation set.

Training GenerRNA on longer sequences enables it to learn more complex patterns, but it also significantly increases computational demands. This is due to the quadratic rise in computational complexity with sequence length in the attention mechanism, coupled with greater memory required for the storage of intermediate results[20]. Additionally, GenerRNA trained on lengthy sequences occasionally generates excessively long outputs, necessitating their truncation. Current computational biology methods struggle with modeling overly long RNA sequences, such as when predicting secondary structures, where there is a challenging trade-off between computational time and accuracy. Such challenges complicate our evaluation efforts. Therefore, we have filtered the pre-training data based on sequence length.

### 2.3 Training Tokenizers

Neural networks are inherently incapable of directly manipulating character-based data, necessitating the conversion of sequences into a tokenized format. In the pre-training phase of our model, we utilize Byte-Pair Encoding (BPE) for this tokenization process[21] as the tokenization strategy. Originally devised as a text compression algorithm, BPE facilitates the representation of variable-length subwords within a fixed vocabulary size. Such capability is notably absent in the k-mer tokenization, which is commonly employed in biological sequence analysis. Additionally, Compared to the one-hot tokenization, where each base corresponds to one token, the BPE tokenizer compresses the information content, enabling the learning and generation of longer sequences. The straightforwardness and effectiveness of BPE tokenization have garnered widespread popularity in the realm of Large Language Models[22].

To train our tokenizer, we constructed a dataset comprising 1 million sequences randomly extracted from RNAcentral. Throughout this procedure, we experimented with various vocabulary sizes ranging from 512 to 50,296 and observed that larger vocabulary sizes led to single tokens encompassing more nucleotides (Supplementary Fig.S1). We opted for a vocabulary size of 1024 primarily for the following two reasons:

1. Drawing on the Chinchilla Scaling Law[23], we calculated the optimal number of tokens considering our computational resources. This estimation guided us in determining the suitable number of nucleotides to include in each token, with the aim of compressing the training data to an ideal number of tokens.
2. In RNA, the chemical bonds at the **nucleotide level** are pivotal in its secondary structure, which subsequently determines its function. Incorporating an overly large number of nucleotides into a single token might potentially overlook this crucial base-level interaction.

### 2.4 Statistical Examination of Nucleotide Propensities

Our research aimed to generate RNA sequences that share similar properties with natural sequences. To realize this, we examined nucleotide distributions under various sampling strategies and parameters. We focused our analysis on three main sampling strategies: greedy search, beam search, and random sampling.

Greedy search persistently chooses the nucleotide with the highest probability, often leading to repetitive sequences. Beam search aims to mitigate this issue by retaining a set of highly probable token sequences within a predefined beam width. We evaluated beam sizes ranging from 5 to 100, incrementing by 5. Owing to the inherent absence of randomness in the outputs under both greedy and beam searches, we initiated each sequence with a distinct permutation of the initial five nucleotides (A, U, G, C). This approach allowed us to generate 1,024 sequences for each parameter configuration in both search methods.

Random sampling chooses from the top k tokens according to the next token’s probability distribution. For this method, we set the temperature at 1.0, a parameter that modulates generation randomness by modifying the logits before the softmax function. We varied the *top*_*k*_ value, ranging from 5 to 1000, creating 1000 sequences for each configuration.

We calculated and compared the distribution of k-mer frequencies between the generated and natural sequences. Our approach utilized a sliding window size of 1bp and a k-value of 3, accommodating a total of 64 unique 3-mers. To quantify the differences in k-mer frequency between the generated and natural sequences, we employed the Kullback-Leibler Divergence (KLD):

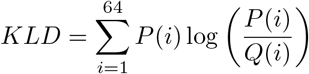

Here, *P* (*i*) represents the frequency of the *i*-th 3-mer in the generated sequences, whilst *Q*(*i*) denotes its frequency in natural sequences. This formula sums over all 64 possible 3-mers(AAA, AAU, …).

### 2.5 Evaluation of Generated RNAs

We investigated the Minimum Free Energy (MFE) of generated RNA sequences as a matrix of secondary structural stability. For the control experiment depicted in Fig.3**a**, we selected 2,000 sequences generated by our model as the **Generated Group**. To lessen the influence of sequence length on MFE, we randomly sampled 2,000 natural sequences with a length distribution identical to that of the generated sequences, forming the **Natural Group**. Notably, longer RNA sequences typically have more opportunities for intramolecular interactions, potentially resulting in lower MFE values[24]. Therefore, by matching the lengths of the sequences in the Generated and Natural Groups, we aimed to make a more meaningful comparison of their respective MFEs. For each sequence in the Generated Group, we created a corresponding sequence of equal length composed of random AUGC nucleotides, designated as the **Random Group**. Additionally, to regulate the effect of nucleotide composition (e.g., GC content), we generated a **Shuffled Group** by thoroughly randomizing the nucleotide order of these sequences. The MFE calculations were performed using the RNAFold[25].

**Figure 2:**
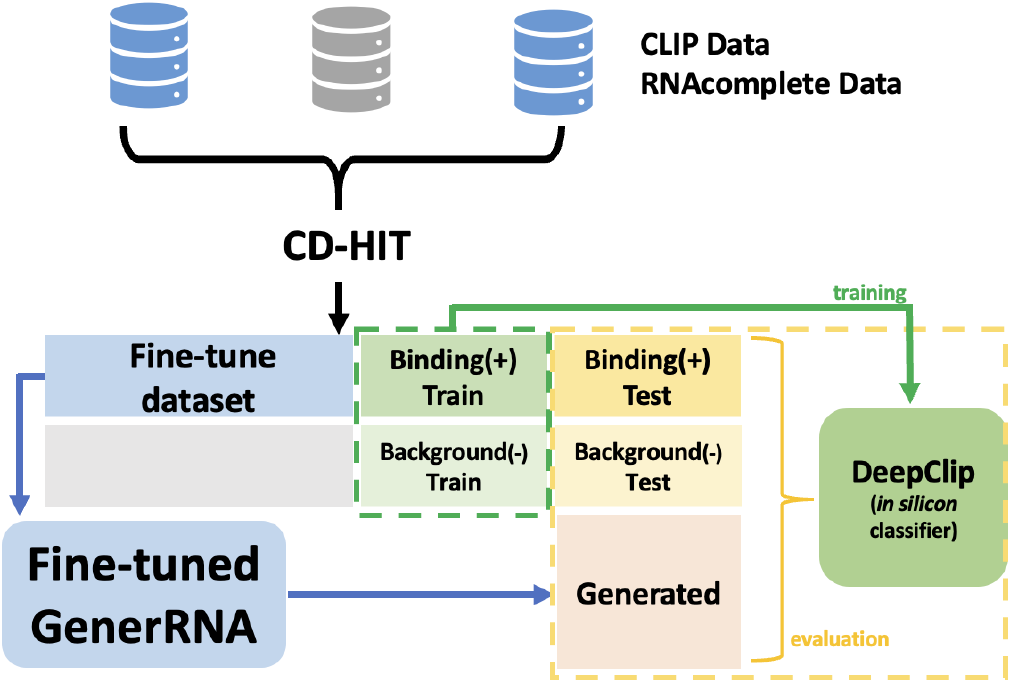
Schematic representation of fine-tuning on protein-binding RNA: The experimental data collected were consolidated and deduplicated using CD-HIT to prevent sequence overlap and data leakage across different datasets. The data were divided into three parts: one for training DeepClip, another for the test set, and the last for fine-tuning the GenerRNA dataset. The trained DeepClip model was subsequently utilized to score sequences generated by GenerRNA.

**Figure 3:**
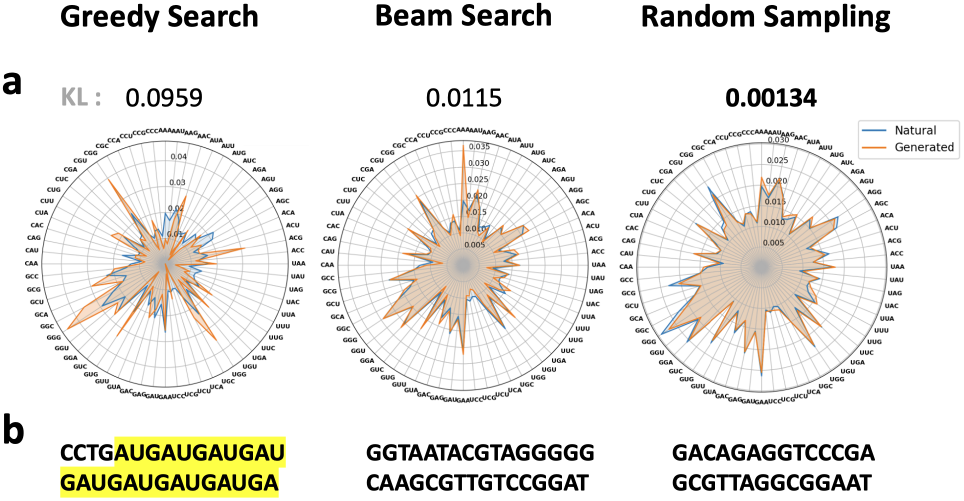
**a**. Radar charts depicting the 3-mer nucleotide distribution of sequences generated under different sampling strategies, accompanied by the Kullback-Leibler (KL) distance comparing these distributions to those of natural sequences. **b**. Examples of generated sequences. **a. b**. Greedy search tends to generate RNA with repetitive fragments, as highlighted, and its 3-mer distribution is significantly different from that of natural sequences (KL distance = 0.0959). Beam search partially alleviates this issue. However, the nucleotide distribution in its generated RNA still deviates from that observed in natural sequences (KL distance = 0.0115). The nucleotide distribution in sequences under random sampling closely aligns with that of natural sequences (KL distance = 0.00134).

Furthermore, to assess the consistency of our model’s performance across various length ranges, we calculated the MFE distribution for sequences of different lengths. We respectively sampled 50 sequences for each length interval, ranging from 10 to 1,000 nucleotides in increments of 10 nucleotides, from the Generated Group. For each length window, we also prepared a corresponding Shuffled Group (Fig.3**b**).

To evaluate the similarity between the generated RNA and known RNA sequences, we employed nhmmer for homology searches[26]. The nhmmer, grounded in hidden Markov models, has been showcased to have superior sensitivity compared to BLAST[27]. We designated all sequences from RNAcentral as the database, with 1,000 generated sequences serving as the query. The search parameters were aligned with those used in the backend of RNAcentral’s sequence search function, and particular details can be found in Supplementary Table.S1.

### 2.6 Fine-tuning GenerRNA on Protein-binding RNAs

With the goal of generating novel RNA sequences that bind to specific proteins, we fine-tuned GenerRNA (Fig.2). We then evaluated the binding affinity between artificially generated RNAs and the target proteins by employing DeepClip[28], an *in silico* method leveraging deep learning techniques to model protein-RNA interactions.

Four *in vivo* CLIP datasets[29, 30, 31] and five *in vitro* RNAcompete datasets[32] were collected for the ELAVL1 protein. ELAVL1 has been implicated in a variety of biological processes and has been linked to a number of diseases, including cancer[33]. The positive sequences (protein-binding sequences) and negative sequences (background sequences sampled from non-target regions) were grouped independently. To minimize the risk of data leakage due to sequence overlap between training and testing sets, we applied a CD-HIT identity threshold of 90% for deduplication. This process yielded 91,136 positive and 106,712 negative sequences. Similarly, for the SRSF1 which act as an oncoprotein and an important target for cancer therapy[34], we obtained 134,343 positive and 137,021 negative sequences from three CLIP datasets[35, 36, 37, 38] and five RNAcompete datasets[39, 32].

Regarding each protein’s binding RNA, 60% of the sequences were allocated for the fine-tuning of GenerRNA, while 30% were used as positive data for training DeepClip, respectively. The remaining 10% was set aside for testing. Subsequently, an equivalent count of sequences from the negative set was randomly selected to function as the negative training, validation, and testing set for DeepClip.

We fine-tuned GenerRNA using the ELAVL1 and SRSF1 Clip datasets, carrying out 50,000 iterations each. The learning parameters were kept consistent with those of the pre-training phase, except for a learning rate of 1e-4 without any warmup or scheduler. Throughout the fine-tuning process, both the validation loss and perplexity exhibited a consistent reduction during the training period.

### 2.7 Evaluation of the Generated Protein-binding RNAs

We employed a fine-tuned version of GenerRNA to generate protein-binding RNAs *de novo*. The decoding parameters used during sequence generation were consistent with those applied in our pre-tuning experiments. The number of sequences generated was equivalent to the number of sequences in our curated test set.

We trained DeepClip over ten epochs, selecting the checkpoint that demonstrated the best performance on the validation set. We then computed binding affinity scores for three distinct sequence datasets: **1. Dataset Generated by GenerRNA, 2. Positive (binding) testset**, and **3. Negative (background) testset**. These affinity scores ranged from a minimum of 0 up to a maximum of 1.0, where a higher score indicates a stronger binding affinity. It should be emphasized that for each specific target protein, the fine-tuning and evaluation of GenerRNA were conducted independently, ensuring tailored optimization for each protein-binding RNA.

### 2.8 Ablation Study

Typically, large-scale model pre-training enhances a model’s generalization performance and robustness. It enables us to fine-tune the model with fewer computational resources and reduced data, facilitating the transfer of acquired knowledge to specific subtasks.

Here, we conducted ablation studies to assess the impact of large-scale pre-training in the context of RNA sequence generation. We trained a model from scratch on ELAVL1-binding RNAs, intentionally bypassing large-scale pre-training, to serve as a control in the ablation group. The utilized model architecture was identical to that in previous experiments. However, we anticipated rapid overfitting due to the smaller volume of training data relative to the model parameters. To address this, we explored the learning parameters, monitored the validation loss, and implemented an early-stopping strategy. Finally, the experiment proceeded through 1500 iterations with a global batch size of 128, equivalent to approximately two training epochs. The ablation model exhibited a slightly higher loss on the validation set compared to the fine-tuned GenerRNA, recording values of 3.87 versus 3.68, respectively. Similarly, the ablation model trained on SRSF1-binding RNAs reached its optimal validation loss during the 750th iteration, with a validation loss of 4.67 which is marginally higher than the 4.60 achieved by the fine-tuned GenerRNA.

Subsequently, we utilized the ablation model and the fine-tuned GenerRNA to independently generate 1000 sequences each. These sequences were then subjected to homology searches using nhmmer, targeting a database containing RNA sequences that bind to target protein. We intentionally set a relatively relaxed threshold to enhance the sensitivity for alignments, and comprehensive details are available in the Supplementary Information.

## 3 Results

### 3.1 Sampling Strategies

Various natural language processing tasks exhibit a propensity for distinct sampling strategies. By measuring the divergence in nucleotide distribution between generated and natural sequences, we identified the most suitable sampling strategies and parameters for generating natural-like RNAs.

The greedy search method opts for the token of the highest probability, leading to consistent results. This approach is particularly effective for structured QA tasks but often results in repetitive segments during long text generation[40]. Notably, a similar phenomenon has also been observed in the generation of RNA sequences. Beam search aims to mitigate this issue by retaining a set of most probable token sequences, proving to be beneficial in tasks needing a balance between efficiency and quality like machine translation[41, 42]. Our results indicate that while RNA sequences generated using beam search tend to exhibit fewer repetitive segments relative to those generated via greedy search, a significant disparity still remains in the distribution of 3-mers between the generated sequences and natural RNA sequences.

Random sampling selects from the top k tokens based on the next token’s probability distribution. This method has been found to output with more randomness and a more natural feel in the long text generation[43]. Our statistical evaluations reveal that the RNA sequences generated using this method exhibit a 3-mer distribution that is closest to that of natural sequences among all assessed sampling strategies.

### 3.2 Minimum Free Energy of Generated RNAs are Similar to that of Natural Ones

The Minimum Free Energy (MFE) of an RNA sequence is defined as the energy of its secondary structure that contributes the least to the free energy, with a lower MFE that signifies a more stable RNA secondary structure.

To eliminate the impact of sequence length on MFE, we ensured a consistent sequence length distribution across all control groups. Additionally, nucleotide distribution in the Shuffled Group was aligned with that of the Generated Group, thereby controlling for factors such as GC content that could affect MFE.

Results indicate that the mean MFE of the generated sequences was slightly lower than that of the natural sequences (-119.1 vs -117.9 kcal/mol), while there was no statistically significant difference between the MFE of two groups (two-sided Mann-Whitney *U* test, p-value = 0.591). Additionally, the MFE of the generated sequences was significantly lower than that of both the shuffled and random groups, as determined by a one-sided Mann-Whitney *U* test (p-value = 3e-06). Besides, Fig.4**c**. depicts that the majority of the generated sequences, regardless of their length, had a lower MFE than its shuffled sequences. These findings indicate that our model can generate sequences with relatively stable secondary structures across a broad range of sequence lengths.

**Figure 4:**
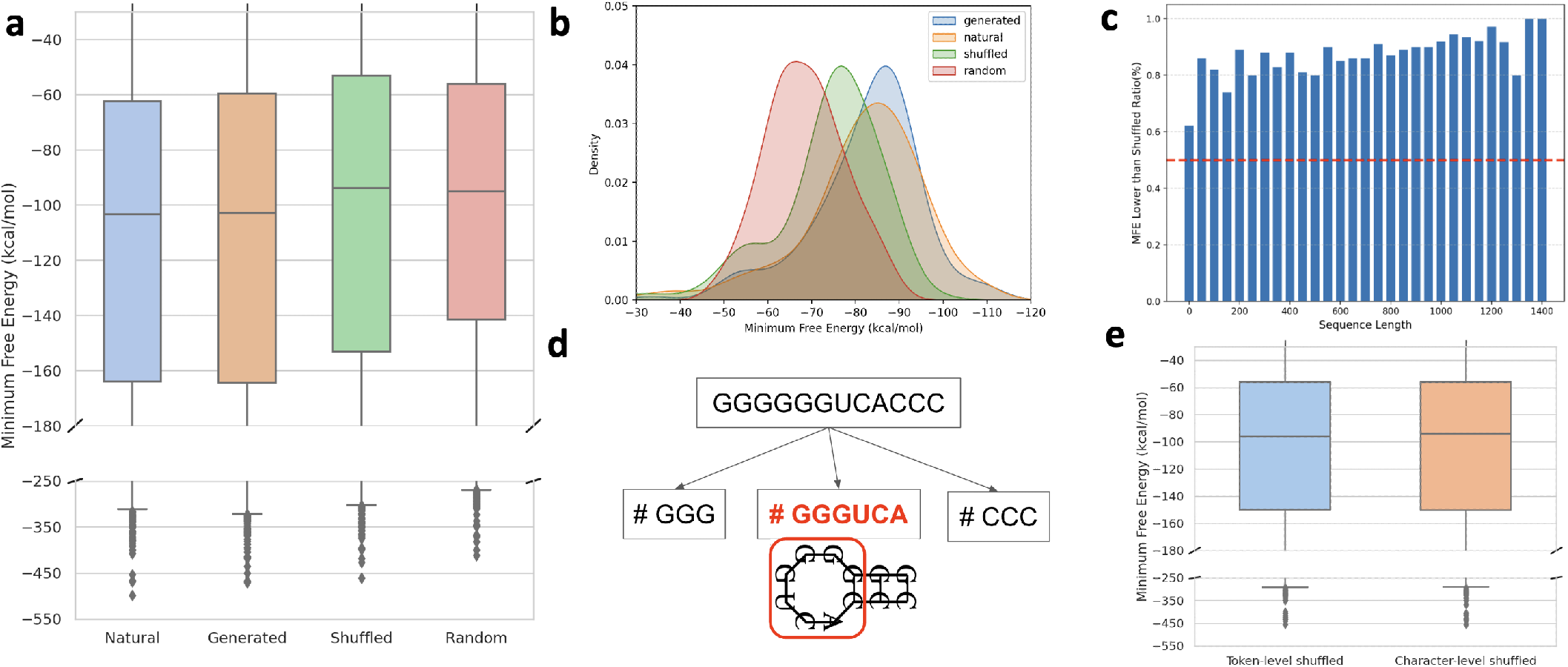
Statistical Analysis of Generated Sequences **a**. This plot displays the distribution of minimum free energy among four groups: Natural Group, Generated Group, Shuffled Group, and Random Group. There is no significant difference in the MFE distribution between generated and natural sequences, whereas the MFE of generated sequences is significantly lower than that of shuffled and random sequences. **b**. MFE distribution among different collections with sequences length set at 250 nt. **c**. Proportion of generated sequences that exhibit a lower MFE compared to their respective shuffled sequences at diverse length intervals. The proportion across all intervals is greater than 0.5, indicating that GenerRNA is able to generate sequences with lower MFE across a variety of length ranges. **d**. Since the BPE tokenizer learns common nucleotide combinations in RNA language, it is possible that it unintentionally learns significant RNA motifs. **e**. When the token sequences generated by the model are shuffled prior to decoding into nucleotide sequences, the MFE distribution of these sequences showed no statistically significant difference compared to the sequences shuffled at the character level. This indicates that the doubt raised in **d** does not hold, or at least does not have a significant impact.

Potential concerns arise given that the Byte Pair Encoding (BPE) tokenizer was trained on a dataset of one million sequences, possibly leading it to learn common nucleotide arrangements and merge these nucleotide combinations into individual tokens. This could inadvertently result in the learning of RNA motifs, which are sequence and secondary structure-related feature patterns in RNA[44]. To investigate whether this was a contributing factor to the observed lower MFEs, we conducted an experiment where the token sequences output by the model were shuffled prior to being decoded back into nucleotide sequences. In this approach, while the nucleotide combinations within the same token stayed adjacent post-decoding, the overall global context was entirely disrupted. We generated 1,000 sequences in this manner, designated as the Token-level-shuffled Group. For comparison, we further shuffled these sequences completely, maintaining the length distribution and base composition unchanged, and denoted these as the Character-level-shuffled Group.

As shown in Fig.4**e**., there was no statistically significant difference in the MFE distribution between these two groups (two-sided Mann-Whitney *U* test, p-value = 0.737). This outcome dispels our initial concerns and substantiates that the lower MFEs of the generated sequences are primarily attributed to the contextual information learned by GenerRNA rather than the BPE tokenizer having learned inherent nucleotide patterns.

### 3.3 Generated RNAs are Distanced from Existing Sequences

To assess the novelty of the sequences generated, a homology search was conducted against a comprehensive database of known sequences. The metric employed for this evaluation was identity, defined as the proportion of matching positions in the aligned subsequences. Additionally, an e-value of 0.1 was set as the threshold for determining the reliability of the alignments.

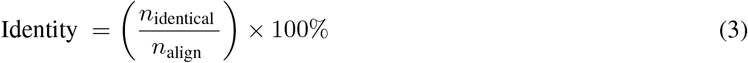

Our findings revealed that 3.6% of the analyzed sequences exhibited 100% identity with those in the database, with an e-value less than 0.1, signifying a perfect match of all nucleotides in the aligned subsequences for these cases. Nonetheless, another 71.5% of the generated sequences aligned with known sequences, meanwhile the aligned subsequences were not identical. Importantly, 24.9% of the sequences did not align significantly with any known sequences at an e-value threshold of 0.1.

As demonstrated in Fig.6**a**., in cases where sequences did not show substantial alignments or had less than 90% identity, the majority still exhibited lower compared to their shuffled ones. This suggests that GenerRNA generates novel RNA sequences that maintain a significant level of secondary structure, even when diverging from known sequences.

**Figure 5:**
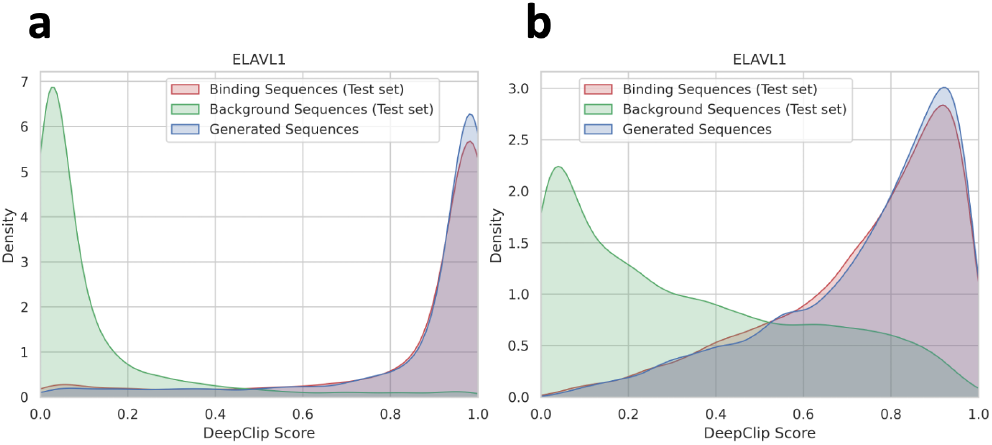
**a.&b**. illustrate the affinity scores of the sequences generated by GenerRNA, along with the positive and negative test sequences, in relation to their respective target proteins (ELAVL1, SRSF1). The affinity scores of the sequences produced by GenerRNA were significantly higher than those of the negative control group, and were comparable to the positive group.

**Figure 6:**
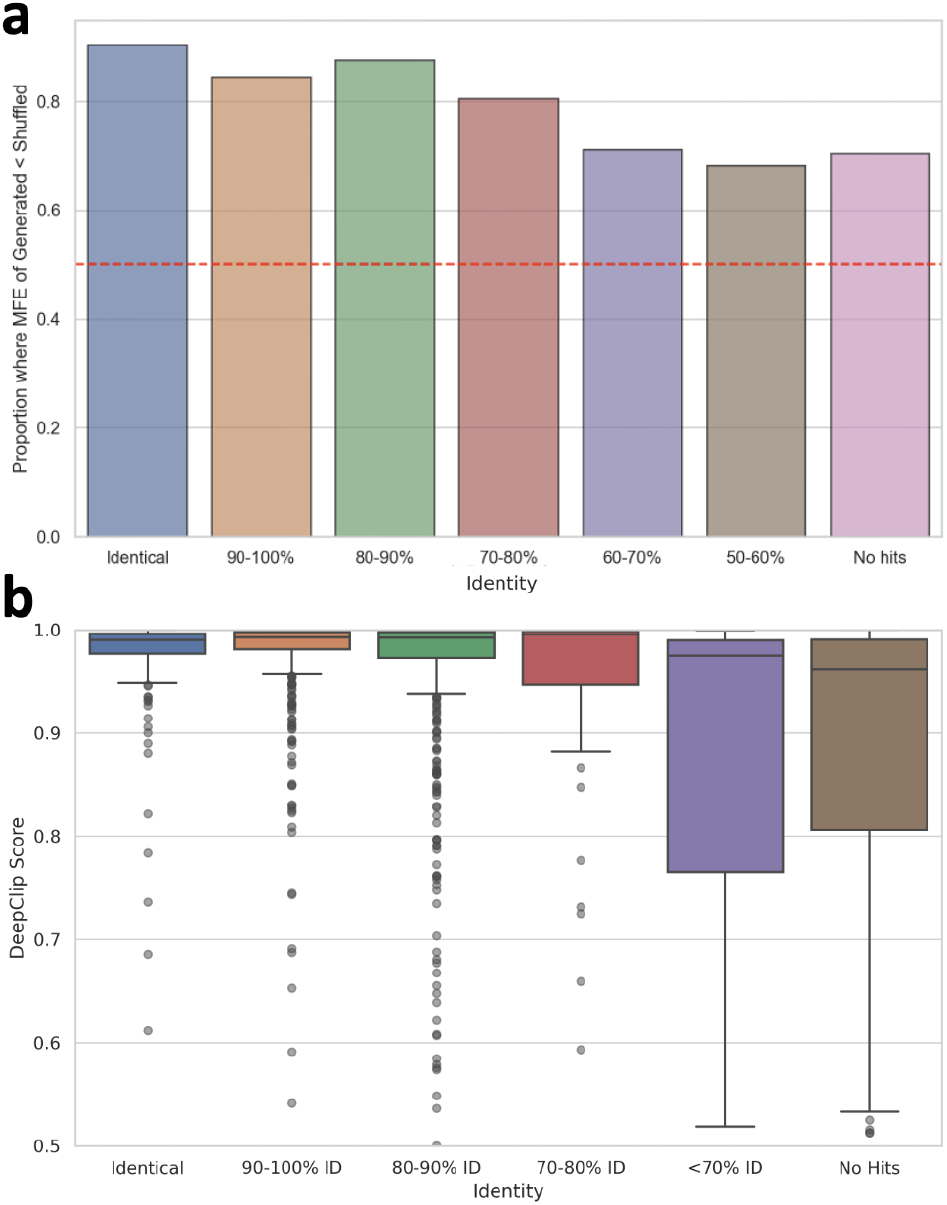
**a**. The ratio of sequences exhibiting lower MFE compared to their corresponding shuffled sequences at diverse identity levels. **b**. Distribution of affinity scores with the target protein (ELAVL1) at varying identity levels. **a.&b**. It is observed that numerous instances of RNA with stable secondary structures or high affinity scores can be found even among sequences with an identity of less than 90% or those that are non-hits.

### 3.4 Fine-tuned GenerRNA generates Protein-Binding RNAs

In the field of natural language processing, the fine-tuning of a pre-trained model on specific tasks boosts task-specific performance or generates articles in a particular style while retaining a general understanding of language syntax, grammar, and semantics. Such a procedure necessitates relatively fewer data and computational resources. In our study, we fine-tuned the GenerRNA using RNA data binding to specific proteins, and estimated the affinity scores between the generated sequences and target proteins using in silico computational methods.

As depicted in Fig.5, sequences generated by GenerRNA exhibit a pronounced specificity toward the target protein. For both examined proteins, the affinity scores of the generated sequences not only significantly exceed those of the negative test set but also marginally surpass the mean scores observed in the positive test set. Specifically, for ELAVL1, the affinity score of generated RNAs was 0.874 compared to 0.858 in the positive test set, and for SRSF1, it was 0.748 versus 0.739.

Furthermore, homology searches reveal that only a small fraction of the RNA sequences generated by GenerRNA that bind to ELAVL1 and SRSF1 proteins were identical to known binding sequences, with 3.12% for ELAVL1 and 1.64% for SRSF1. Notably, a majority of the sequences, 76.0% for ELAVL1 and 69.6% for SRSF1, was not aligned with any known binding sequences. As illustrated in Fig.6**b**., many instances of RNA with high affinity scores were observed even among sequences with less than 90% identity or those that did not form alignments. It is important to mention that the tool adopted for our homology analyses is highly sensitive, and we have intentionally set more relaxed thresholds to bolster the detection sensitivity for alignments.

Moreover, we conducted an ablation study of the impact exerted by large-scale pre-training on tasks related to the generation of RNA sequences in downstream applications. Our investigation involved a comparasion between a ablation model devoid of pre-training, and the GenerRNA undergone both pre-training and fine-tuning. The ablation model, despite exhibiting a slighter higher but close validation loss compared to GenerRNA, could only produce sequences characterized by a lack of originality. These sequences predominantly mirrored existing data, indicating a tendency towards mere replication rather than genuine generation. We speculate that the pre-training on extensive datasets typically empowers the model to internalize a wide array of features and patterns inherent to the RNA data, which offers a robust starting point that enhances the model’s capacity for fine-tuning on specific tasks. This leads to improved generalization and the ability to generate RNA sequences that are not only novel but also adhere to the “grammar” of RNA.

**Table 1:**
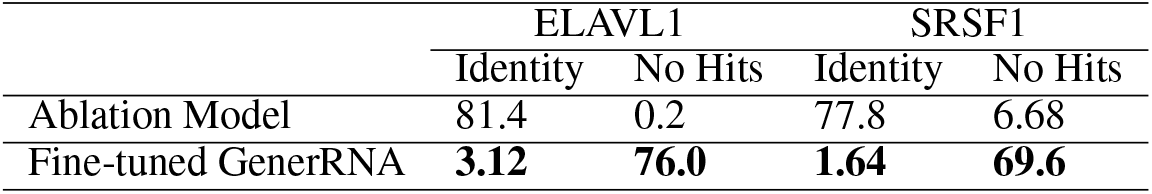
Comparison of sequence novelty in ablation model and fine-tuned GenerRNA: the fine-tuned GenerRNA generates a greater number of novel RNAs unaligned with known RNAs and fewer sequences identical to known RNAs.

|  | ELAVL1 |  | SRSF1 |  |
| --- | --- | --- | --- | --- |
|  | Identity | No Hits | Identity | No Hits |
| Ablation Model | 81.4 | 0.2 | 77.8 | 6.68 |
| Fine-tuned GenerRNA | **3.12** | **76.0** | **1.64** | **69.6** |

## 4 Discussion

Biological sequences harbor a wealth of information, covering evolutionary history, survival strategies, and even blueprints for future development. AI language models serve as translators for reading and writing this enigmatic language. The ever-increasing number of RNA sequences in public databases, coupled with advancements in natural language model architectures, provides a foundation for constructing models proficient in generating biological sequences.

Our development of GenerRNA marks the first instance of a large-scale, pre-trained RNA generation model. Trained on 17.4 billion nucleotides, it learns the grammar and vocabulary of the RNA language and is capable of “speaking” this language. GenerRNA is able to generate RNA that possesses secondary structural stability akin to known sequences while preserving a distinctiveness from them. Our approach is devoid of reliance on predefined secondary structures or any ancillary prior knowledge, aligning with a zero-shot approach. This zero-shot style generation holds the potential to expand the known RNA space, enriching our understanding of RNA structures and functions.

In our study, we elucidated that a fine-tuned GenerRNA can generate RNA sequences with high binding affinity to target proteins while maintaining uniqueness from existing sequences. This approach not only represents a groundbreaking and effective method for generating potential pharmaceutical candidates but also demonstrates the practicality of GenerRNA as a highly versatile platform for RNA generation.

GenerRNA represents the initial advancement of large-scale generative language models in the realm of RNA. We anticipate that future developments in RNA generative language models will cover numerous directions yet not be confined to them.

One promising avenue is to explore downstream applications of generative RNA language model. These include generation of functional RNAs, aiding in the design of RNA vaccines, and the development of nucleic acid therapeutics.

Another direction involves scaling. Similar advances in protein-related fields have shown that models with larger parameters size generate novel protein structures more effectively and may demonstrate emergent abilities in specific tasks[45, 46]. This potential scalability could further utilize the ability of the RNA language model, warranting further investigation.

Lastly, controlled generation is also a valuable field to develop. Techniques such as prompt tuning[47] could be instrumental in directing these models to generate sequences that belong to specific families or possess desired characteristics.

