## supplemental for "GenerRNA: A generative pre-trained language model for *de novo* RNA design"

Supplementary Information

Yichong Zhao, Kenta Oono, Hiroki Takizawa, and Masaaki Kotera

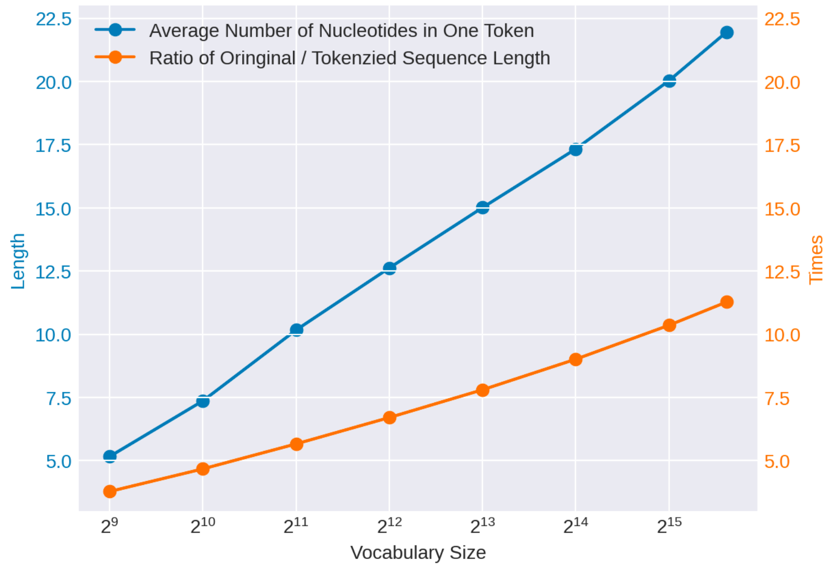


**Fig.S1. Effect of Vocabulary Size on Token Length and Ratio of Original/ Tokenized Sequence Length**

Our training dataset encompassed 17.4 billion nucleotides. Draw on some rules of thumb [1] suggesting that the optimal data size under our computational resources is 3~5 billion tokens, we selected a vocabulary size of 1024 for our BPE tokenizer.

**Table.S1. Hyperparameters of *nhmmer* for homology search**

| | **Query: RNA Generated by Pre-trained Model<br>DB: RNAcentral** | **Query: RNA Generated by Fine-tuned Model<br>DB: Curated Protein-binding RNA** | **Remark** |
|---|---|---|---|
| **-T** | **0** | **0** | Report target sequences with a bit score >= T |
| **--F3** | 0.02 when sequence length < 50 nt | - | A heuristic acceleration parameter |
| **--watson** | True | False | Only search the top strand |
| **-Z** | Number of Sequence in RNAcentral | Number of Sequence in RNAcentral | Size of the target database used for E-value calculation |

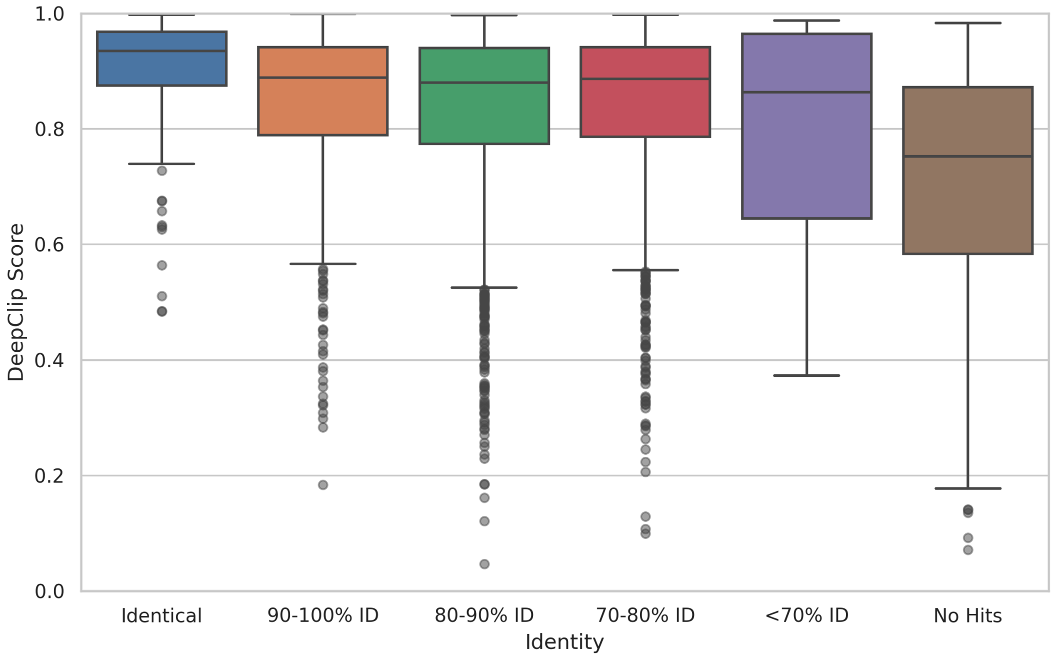


**Fig.S2. Distribution of affinity scores with the target protein (SRSF1) at varying identity intervals.**

In line with the experimental outcomes illustrated in Fig.6.b of the main text, the sequences generated by the fine-tuned GenerRNA encompass numerous RNA instances that, while exhibiting lower identity and, in some cases not aligned with any known sequences, nonetheless possess significantly elevated affinity scores.

### Reference:

1. Jordan Hoffmann, Sebastian Borgeaud, Arthur Mensch, Elena Buchatskaya, Trevor Cai, Eliza Rutherford, Diego de Las Casas, Lisa Anne Hendricks, Johannes Welbl, Aidan Clark, Tom Hennigan, Eric Noland, Katie Millican, George van den Driessche, Bogdan Damoc, Aurelia Guy, Simon Osindero, Karen Simonyan, Erich Elsen, Jack W. Rae, Oriol Vinyals, Laurent Sifre Training compute-optimal large language models. arXiv preprint arXiv:2203.15556, 2022.
